# Peptide-stimulation of autologous T cells reverts immune checkpoint inhibitor resistance of hypermutated colorectal cancer cells

**DOI:** 10.1101/2024.02.06.579061

**Authors:** Sandra Schwarz, Su Zhaoran, Mathias Krohn, Markus W. Löffler, Andreas Schlosser, Michael Linnebacher

**Affiliations:** Molecular Oncology and Immunotherapy, Department of General Surgery, University Medicine Rostock, Rostock, Germany; Department of General, Visceral and Transplant Surgery, University Hospital Tübingen, Tübingen, Germany; Department of Immunology, Interfaculty Institute for Cell Biology, University of Tübingen, Tübingen, Germany; German Cancer Consortium (DKTK) and German Cancer Research Center (DKFZ) Partner Site Tübingen, Tübingen, Germany; Cluster of Excellence iFIT (EXC2180) ‘Image-Guided and Functionally Instructed Tumor Therapies’, University of Tübingen, Tübingen, Germany; Department of Clinical Pharmacology, University Hospital Tübingen, Germany; Rudolf-Virchow Center, Center for Integrative and Translational Bioimaging, University of Würzburg, Würzburg, Germany

## Abstract

Two hypermutated colon cancer cases with patient-derived cell lines, peripheral and tumor-infiltrating T cells available were selected for detailed investigation of immunological response.

T cells co-cultured with autologous tumor cells showed only low levels of pro-inflammatory cytokines and failed at tumor recognition. Similarly, treatment of co-cultures with immune checkpoint inhibitors (ICI) did not boost antitumor immune responses. Since proteinase inhibitor 9 (PI-9) was detected in tumor cells, a specific inhibitor (PI-9i) was used in addition to ICI in T cell cytotoxicity testing. However, only pre-stimulation with tumor-specific peptides (cryptic and neoantigenic) significantly increased recognition and elimination of tumor cells by T cells independently of ICI or PI-9i.

We showed, that ICI resistant tumor cells can be targeted by tumor-primed T cells and also demonstrated the superiority of tumor-naïve peripheral blood T cells compared to highly exhausted tumor-infiltrating T cells. Future precision immunotherapeutic approaches should include multimodal strategies to successfully induce durable anti-tumor immune responses.

## 1. Introduction

The interaction of tumor with immune cells triggers dynamic adaption to evade immunosurveillance. A prominent example is tumors’ genetic or epigenetic modifications to reduce or switch-off HLA class I expression, thus preventing detection by T cells. Depending on the tumor entity, up to 80% of solid tumors have completely lost HLA class I expression ^1^. Another tool to repress antitumor immune responses is the expression of immunomodulatory proteins like programmed cell death protein 1 (PD-1), cytotoxic T lymphocyte associated protein 4 (CTLA-4) and lymphocyte activation gene 3 (LAG-3) or their respective ligands in the tumor microenvironment ^2^. Via these immune checkpoint molecules, tumor cells utilize the physiologically programmed down-modulation of activated immune cells. However, therapeutic interfering with these immunosuppressive mechanisms via immune checkpoint inhibitors (ICI) represents the latest revolution in cancer treatment.

The application of ICI is mostly limited to tumors with high mutational burden (TMB), since clinical trials proved highest response rates in this patient group (reviewed in ^3^). However, despite preselection of patients, studies with microsatellite instable colorectal cancer (CRC) revealed, that up to 70% of patients do not benefit from pembrolizumab ^4^, nivolumab ^5^ or durvalumab ^6^ treatment. Even though combining ipilimumab and nivolumab raised patients’ objective response rates to 55% ^7^, still a substantial number of hyper-mutated tumors resist ICI treatment.

High TMB and constitutive HLA class I expression are both favorable prognostic markers for immunotherapy ^8,9^, but ICI therapy response prediction remains unsatisfactory. High TMB of ≥10 mutations/Mb is insufficient to predict ICI success, neither across cancer entities nor within a specific one ^10,11^. However, the combination of TMB and loss of heterozygosity in HLA class I improved differentiation between ICI therapy responders and non-responders in lung cancer ^12,13^. Similarly, a score retrospectively determining tumor immunogenicity by TMB combined with antigen processing machinery gene expression improved ICI therapy response prediction in urothelial cancer and melanoma datasets ^14^.

T cells interaction with tumors can further be impaired by several other immunosuppressive mechanisms beyond dysfunctional antigen presentation. Malignant cells often secrete immune regulatory molecules, *e.g.* cytokines, which exert immunosuppressive effects in the tumor microenvironment ^15,16^. Likewise, enzymes and enzyme inhibitors with a role in regulating immune cell activity have been identified as complementary tumor immune evasion strategies ^17,18^.

Here, we selected two microsatellite instable CRC cases exhibiting constitutive HLA expression and investigated the interaction of T cells with autologous tumor cells *ex viv*o. Co-culture of tumor cells with autologous peripheral (pTc) or tumor-infiltrating T cells (TiTc) was applied to test the antitumor effect of isolated T cells alone or in combination with immune modulatory agents matching tumor cells’ characteristics. Moreover, effects of individualized inhibition of immunosuppression were directly compared to anti-tumor immune responses of autologous T cells recognizing tumor-specific antigenic peptides.

## 2. Results

### 2.1 Selection of tumor cases

For a detailed analysis of tumor and T cell interactions, appropriate patient biomaterials were selected from our comprehensive set of CRC models available at the BioBank Rostock ^19^.

Considering all decisive factors (biomaterial availability, frequent non-synonymous mutations, antigen processing, HLA class I expression and HLA class II inducibility), the two CRC cases HROC113 and HROC285 were selected. Both tumors showed high TMB (HROC113: 176 mutations/Mb; HROC285: 212 mutations/Mb) and microsatellite instability (MSI). Tumor cell lines were established directly from the patient’s tumor (HROC113) or from a patient-derived xenograft (HROC285 T0 M2). TiTc and pTc as well as B-LCLs were also available for both patients.

### 2.2 T cells do not get activated by co-culture with autologous tumor cells

Co-culture experiments of T cells and autologous tumor cells were performed to investigate if the tumors with high TMB and HLA expression can induce sufficient immune responses. Tumor cells were incubated with T cells, which were pre-treated as follows: 1) expanded T cells, 2) T cells, which were expanded and co-cultured with autologous tumor cells for 14 days and 3) T cells, which were expanded and cultured for 14 days in the same conditions (temperature, media, supplements, change of media) but without tumor cells (= control culture with tumor-naïve T cells). In the degranulation assay (Figure 1A), the amount of T cells recognizing tumor cells (CD8^+^/CD107a^+^/IFNγ^+^ cells) was larger in co-cultured compared to unspecifically expanded pTc HROC113 (p=0.04). Similarly, more degranulating T cells were detected among tumor-naïve pTc HROC113 from the control culture than among unspecifically expanded pTc HROC113 (p=0.02). There was no significant difference observed between the reactivity of co-cultured and control-cultured pTc HROC113 (p>0.05; Figure 1A). The TiTc HROC113 showed a similar pattern (Figure 1B) and corroborated the previous results (p_co-culture vs expanded Tc_=0.02; p_control-culture vs expanded Tc_=0.03; p_co-culture vs control culture_>0.05). In contrast, the population of tumor-reactive cells among pTc HROC285 was similar in all three tested conditions (unspecific expansion, co-culture, control culture) and no distinct differences were observed (Figure 1C). TiTc HROC285 were also assessed using this experimental approach but unfortunately the proportion of CD8^+^ cells, being as low as 27% in the beginning, even decreased further during the cultivation period and no reliable measurements of CD8^+^/CD107a^+^/IFNγ^+^ cells could be obtained.

**Figure 1:**
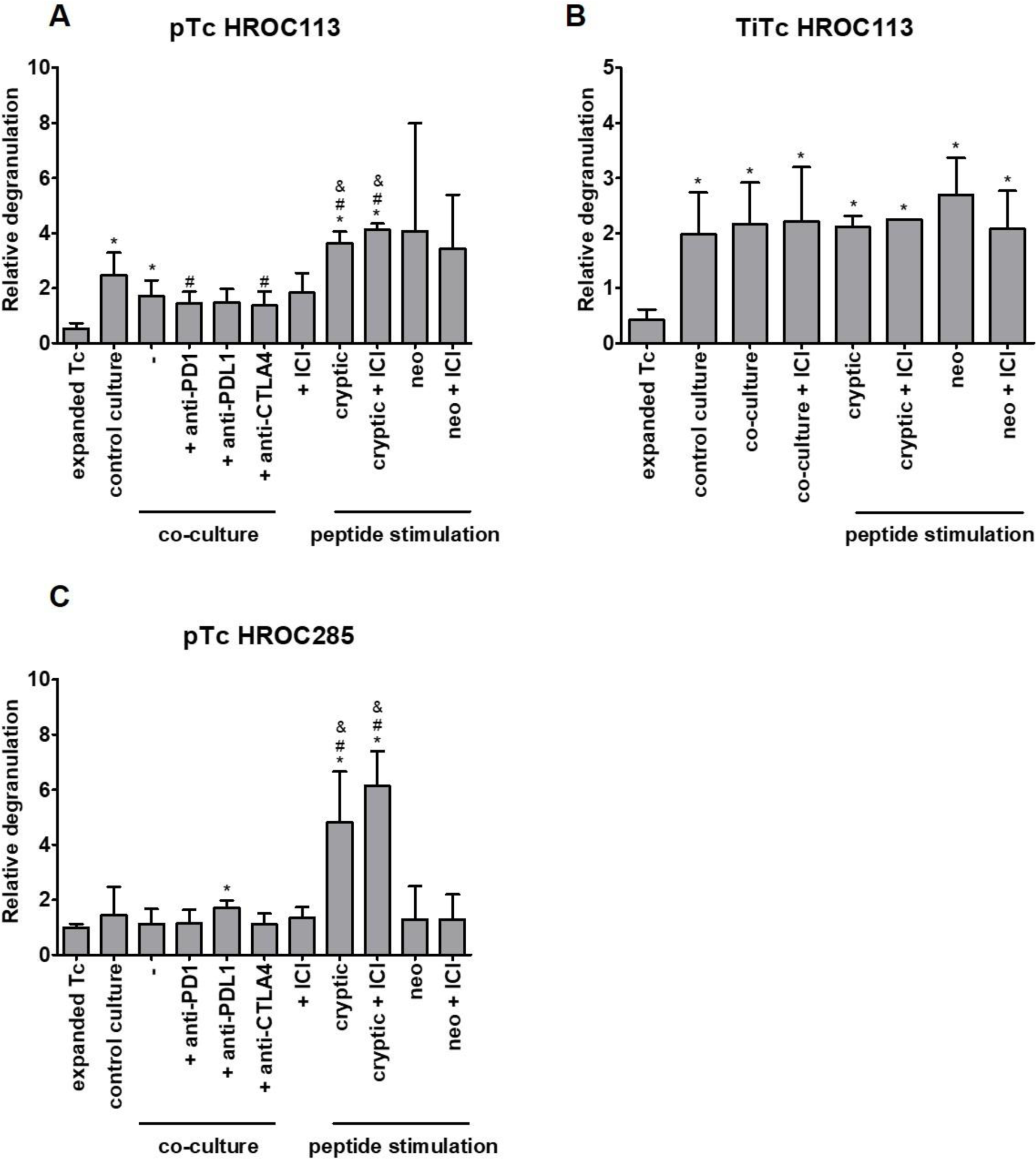
Effect of co-culture, ICI treatment and peptide stimulation on tumor cell recognition by Tc. (A) pTc HROC113, (B) TiTc HROC113 and (C) pTc HROC285 underwent different (co-)culture conditions or were stimulated with tumor-specific peptides to increase their antitumor reactivity measured by degranulation assay after 5h incubation with freshly prepared tumor cells. CD8^+^/CD107a^+^/IFNγ^+^ cells were considered tumor-reactive. Results were normalized to measurements of control Tc not incubated with tumor cells. p<0.05 in *t*-test of sample vs. expanded T cell (*), control culture (#) or co-culture (&). Depicted are means of 2-6 biological replicates and the respective standard deviation.

### 2.3 ICI do not increase the recognition of autologous tumor cells by T cells

Since tumor cells failed to induce T cell responses, ICI (pembrolizumab (anti-PD-1), durvalumab (anti-PD-L1), and ipilimumab (anti-CTLA-4)) were added to the co-culture to improve anti-tumor reactions. Contrasting expectations, pTc HROC113 recognized tumor cells better after control culture than after ICI-supplemented co-culture, showing significantly deterioration after co-culture with PD-1 or CTLA-4 supplementation (p=0.048 and p=0.041, respectively) (Figure 1A). Moreover, the degranulation capacity of co-cultured pTc HROC113, with and without ICI treatment, was significantly lower compared to tumor-naïve pTc HROC113 (p=0.004).

Due to the low quantity of available TiTc HROC113, single ICI testing was replaced by a combination treatment (Figure 1B) but did also not improve tumor cell recognition. The results of pTc HROC285 (Figure 1C) resembled those of pTc HROC113: tumor-naïve pTc HROC285 showed high degranulation, contrasting co-cultured pTc HROC285 with decreased tumor cell recognition. The addition of ICI was insufficient to overcome the tumor cell-induced T cell inhibition.

### 2.4 HROC113 and HROC285 T0 M2 express genes of the antigen-processing machinery

To investigate reasons for the failure of ICI treatment *in vitro*, we analyzed the expression of genes involved in antigen processing and presentation. By determining the expression of 18 genes used in the study of Wang et al ^14^, their immunological functionality was assessed in the established tumor cell lines. Respective results were compared to datasets of normal colon and CRC samples from TCGA and GTEX (Colon: TCGA normal: n=51; TCGA tumor: n=484; GTEX normal: n=308). Surprisingly, when compared to normal colon samples, gene expression was higher in both tumor cell lines as well as in TMB high and TMB low tumor tissues regarding 16/18 genes investigated (Figure 2A-B). Only the expression of β2 microglobulin (B2M) was found significantly lower in tumor tissues compared to healthy colon samples in the analyzed data sets. When assessing differing genes between TMB high and low CRC tissue samples, we observed, that the expression level of ERAP1, PDIA3, PSMB6, PSMB9, PSMB10, CALR, CANX, TAP1, TAP2, TAPBP, B2M and HLA-C was found to be lower in tumors featuring fewer mutations. In both tumor cell lines, the mean expression of the selected genes resembled or even exceeded the levels in CRC samples from TCGA, but due to the small size of the cell line cohort (n=2), statistical analysis was inadmissible. Moreover, functional antigen processing was previously assured through HLA ligandome analyses enabling the characterization of several thousand HLA ligands for each CRC cell line, including tumor-specific antigens ^20^.

**Figure 2:**
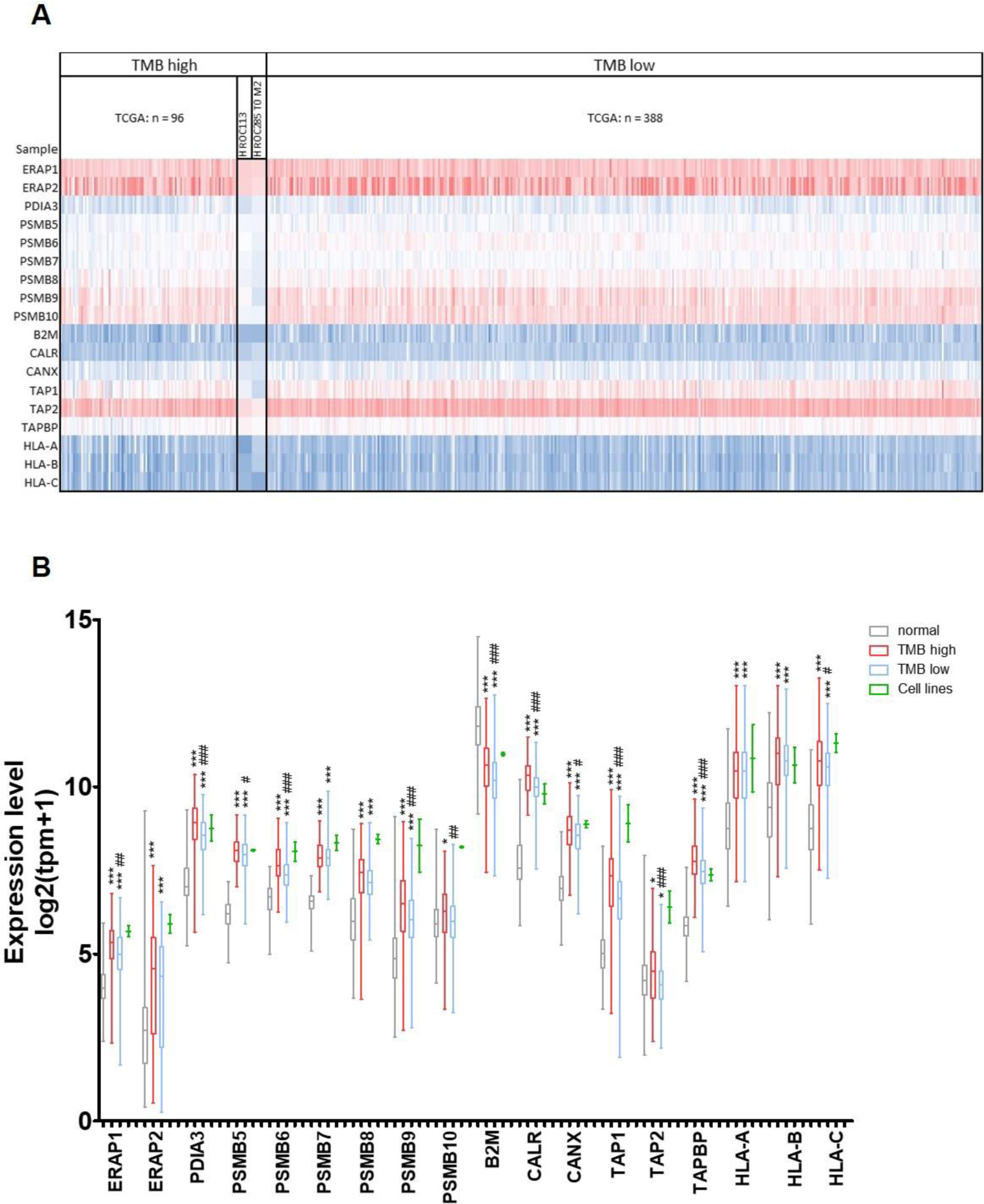
Expression of genes involved in antigen processing. TCGA data of CRC (n=484) were compared with normal colon tissue (TCGA: n=51, GTEX: n=308) and tumor cell lines (n=2), regarding expression of 18 genes involved in antigen processing and presentation. Tumor cell line data contained three technical replicates each, but due to the small group size, statistical calculations were omitted. (A) Heatmap of analyzed data sets depicting high expression in blue and low expression in red, in regard to TMB. (B) Expression of antigen processing genes; *t*-test of tumor sample vs. normal tissue with p<0.05 (*), p<0.01 (**) and p<0.001 (***); *t*-test of TMB high vs. TMB low with p<0.05 (#), p<0.01 (##) and p<0.001 (###).

Flow-cytometry analyses showed that 98% of HROC113 and 96% of HROC285 T0 M2 cells stained positive for HLA class I (Figure 3A). IFNγ pre-treatment increased the amount of HLA class I/II double positive cells in HROC113 from <1% to 45% (p<0.001). In HROC285 T0 M2, the proportion of HLA class I^+^/II^+^ cells even reached 72% following IFNγ treatment compared to 2% in untreated cells (p<0.001). Accordingly, median fluorescence intensity of HLA class II increased by 20-fold after IFNγ stimulation (p=0.002). Thus, IFNγ pre-treatment was performed for all the following experiments.

**Figure 3:**
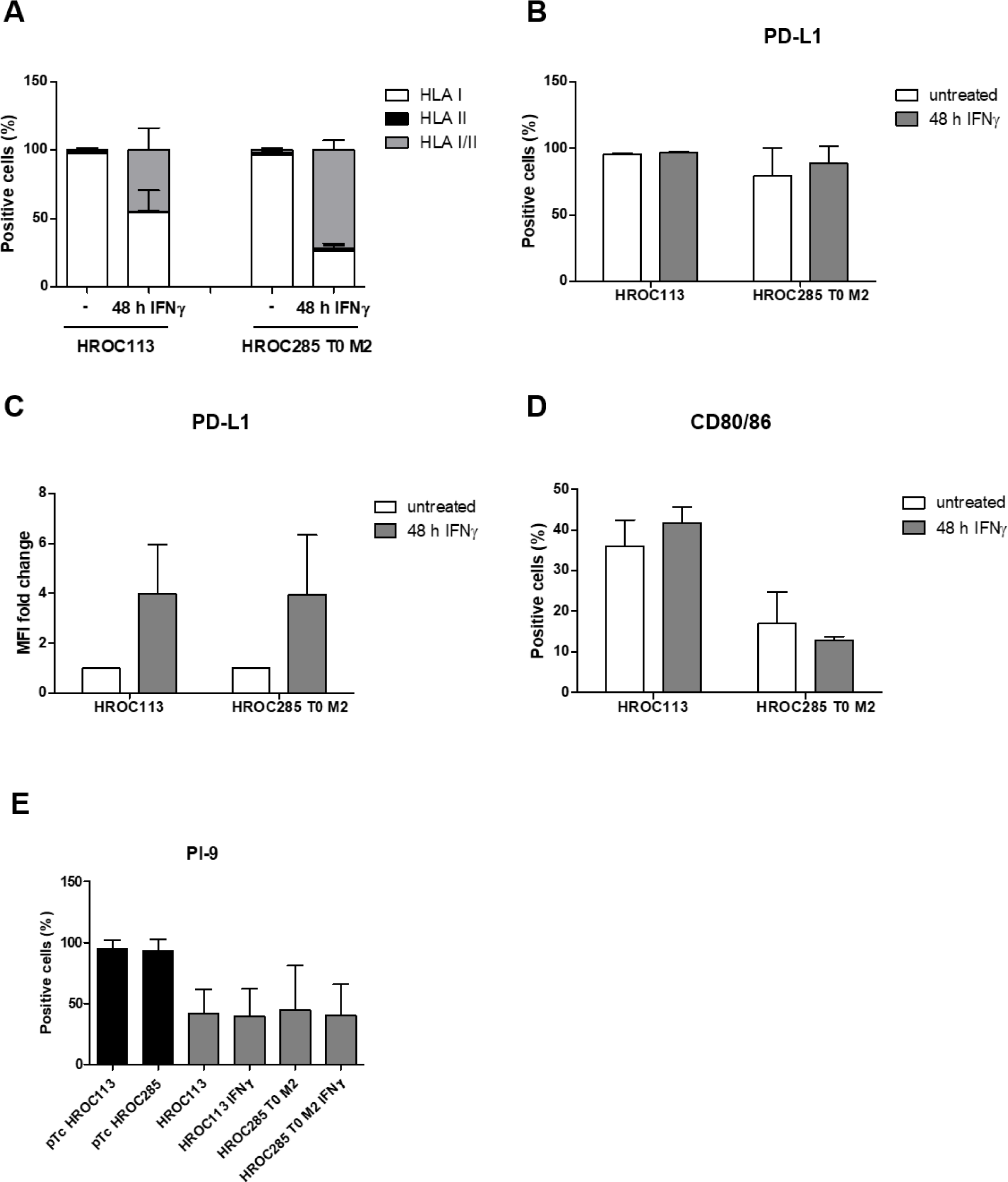
Tumor cell characterization. Untreated or IFNγ-treated (200IU/ml for 48h) tumor cells were analyzed via flow cytometry. (A) shows PD-L1 positive cells. (B) Median fluorescence intensity was determined and fold change in IFNγ-treated and untreated tumor cells was calculated. (C) Percentage of HLA class I^+^ and II^+^, (D) CD80^+^/86^+^ and (E) PI-9^+^ were determined. Depicted are means of 3-5 biological replicates and the respective standard deviation.

These results prove the integrity of antigen presentation in the selected tumor cell lines suggesting functional immune interaction with autologous lymphocytes. Thus, the observed immunosuppression by the tumor cells must be ascribed to other immune evasion mechanisms.

### 2.5 Tumor cells utilize several immunosuppressive mechanisms

Expression of the inhibitory checkpoint molecule programmed death-ligand 1 (PD-L1) was observed in the majority of tumor cells (HROC113: 95%; HROC285 T0 M2: 79%; Figure 3B-C). Here, stimulation with IFNγ increased median fluorescence intensity of PD-L1 expression about 4-fold in both tumor cell lines (p=0.05 in both cases). When determining CD80/86 expression, this CTLA-4 ligand was detected on 30% and 17% of HROC113 and HROC285 T0 M2 cells, respectively (Figure 3D). Treatment with IFNγ did not significantly change CD80/86 expression.

Further characterization of the tumor cells revealed granzyme B activity in HROC113 as well as HROC285 T0 M2, which was found independent of IFNγ treatment (Figure S1). As this apoptosis-propagating enzyme threatens cell survival, its presence is often accompanied by the expression of proteinase inhibitor 9 (PI-9), which specifically inhibits the function of granzyme B. Indeed, at least 40% of HROC113 as well as HROC285 T0 M2 tumor cells expressed this enzyme inhibitor (Figure 3E). This finding seems to be common in CRC cells, since PI-9 expression was also observed in several other patient-derived CRC cell lines in varying amounts and was independent of MSI (Figure S2).

The analysis of secreted cytokines showed that neither HROC113 nor HROC285 T0 M2 produced IL-10 in measurable amounts, which is known to impair T cell function. Additionally, the cytokine profile of the tumor cell lines (Figure S3) did not explain their immunosuppressive behavior in co-culture. Fibrinogen-like protein 1 (Figure S4), an immunosuppressive ligand of LAG-3, was also not detectable in tumor cell culture supernatants.

### 2.6 T cells characterization shows exhausted and regulatory subpopulations

Poor tumor cell recognition can be caused by an impaired ability for activation due to T cell exhaustion. To assess the level of exhaustion in unspecifically expanded pTc and TiTc from patients HROC113 and HROC285, expression levels of PD-1, CTLA-4 and LAG-3 were determined. LAG-3 (pTc HROC113: 82%; pTc HROC285: 67%) and CTLA-4 (pTc HROC113: 68%; pTc HROC285: 51%) were found to be expressed on the majority of pTc, while PD-1 was present on less than 20% of T cells (Figure 4A-B). Exhausted T cells were classified as PD-1/CTLA-4/LAG-3 triple positive cells with 10% of pTc HROC113 and 9% of pTc HROC285 identified as such, respectively.

**Figure 4:**
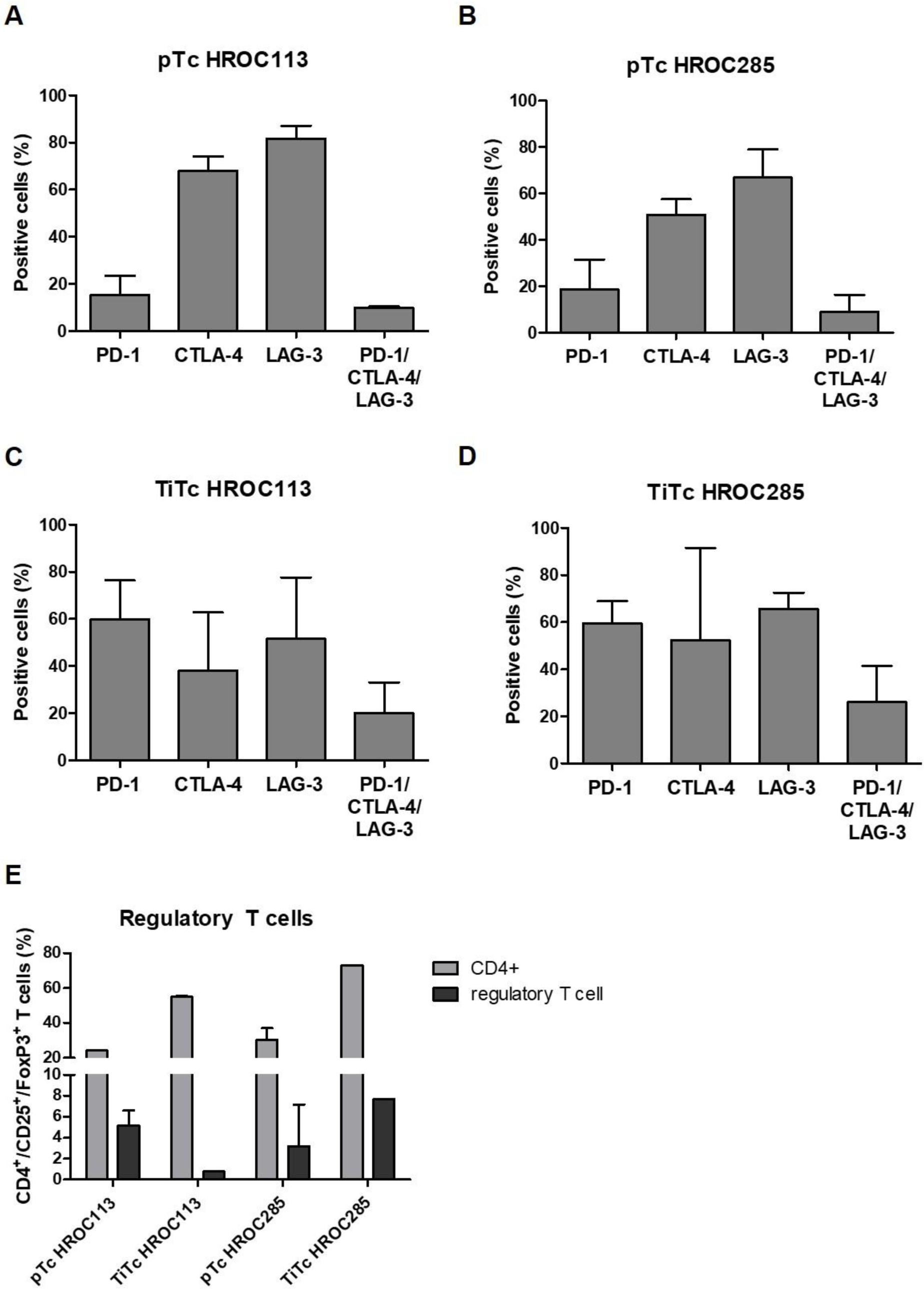
Characterization of T cells. Following 14 days of unspecific expansion, Tc were analyzed regarding (A-D) exhaustion markers and (E) proportion of regulatory Tc defined as CD4^+^/CD25^+^/FoxP3^+^. Depicted are means of 3-4 biological replicates and the respective standard deviation.

The assessment of immune checkpoint molecules on TiTc (Figure 4C-D) revealed a larger quantity of cells positive for PD-1 in TiTc of HROC113 and HROC285 compared to the respective matching pTc (p=0.02). The large proportion of PD-1^+^ TiTc (60% in TiTc HROC113 as well as TiTc HROC285) was accompanied by an overall enlarged population of exhausted cells in TiTc (p=0.05).

Even though, the amount of CD4^+^ cells was higher in TiTc than pTc (p<0.001), this did not coincide with larger amounts of regulatory T cells (Figure 4E). In pTc HROC113, 5.2% of the cells were CD4^+^/CD25^+^/FoxP3^+^, while this population represented less than 1% of cells among the matching TiTc (p=0.006). A contrasting observation was made in pTc and TiTc HROC285, where 3.2 and 7.7% of the cells were identified as regulatory T cells, respectively.

The cytokine detection assay in T cell and co-cultures revealed secretion of the pro-inflammatory interleukins (IL) −5, −6 and −17a as well as GM-CSF (Figure S3). It was frequently seen, that the cytokine concentration was highest in T cell culture and decreased in the co-culture setting, while secretion by tumor cells alone was minimal.

### 2.7 Peptide-stimulated T cells recognize autologous tumor cells better than co-cultured T cells

In a previous study, we showed that T cell stimulation with mutated neoantigens and non-mutated cryptic peptides improved tumor cell recognition in pTc and TiTc ^20^. In the present study, we compared the strength of T cell activation mediated by these tumor-specific peptides with T cell stimulation by autologous tumor cells. pTc HROC113 and pTc HROC285 pre-stimulated with cryptic peptides reached significantly higher amounts of degranulating cells compared to all respective co-culture approaches (p<0.001 in both cases) (Figure 1A and C). A similar effect concerning improved tumor cell recognition by neoantigen stimulation was observed in pTc HROC113, but high variance prevented reaching significance. Additionally, these peptide stimulation experiments showed that tumor-specific T cell responses can be induced similarly in both pTc and TiTc, while tumor cell recognition by pTc partly even proved superior (p_pTc HROC113 (cryptic peptides) vs TiTc HROC113 (cryptic peptides)_=0.01). Furthermore, these results demonstrate, that the tumor cells are indeed immunogenic and the missing T cell response in co-culture tests must be ascribed to remarkable immunosuppressive tumor cell properties. However, since the combined ICI treatment could not improve tumor cell recognition by peptide-stimulated T cells, we assume that for these two MSI CRC cases, immunosuppression does not only depend on PD-1, PD-L1 and CTLA4.

### 2.8 Peptide-stimulated pTc HROC113 eliminate autologous tumor cells

Since granzyme B activity measurement was unsuitable for detecting T cell-mediated tumor cell killing, we set up an *in vitro* cytotoxicity assay based on crystal violet staining of adherent tumor cells. These were stimulated with IFNγ and treated with ICI and/or a PI-9i, which had been proven to successfully control tumor growth in mice by Jiang et al. ^18^. The effect of the PI-9i on tumor and T cell viability was tested beforehand with concentrations covering 25-1,000µM (Figure S5). The highest concentration not inducing cytotoxic effects, i.e. 400µM, was chosen for application in the cytotoxicity assay.

Cytotoxicity was experimentally tested as a proof of concept with peptide-stimulated pTc HROC113, as they showed significantly better tumor cell recognition than co-cultured T cells. After 48h of incubation with autologous tumor cells, the cytotoxic capacity of untreated control pTc HROC113 was low (3% tumor cell killing), and was not significantly improved by the addition of ICI, PI-9i or the combination of both (Figure 5). On the contrary, pTc HROC113 pre-stimulated with tumor-specific cryptic peptides or neoantigens eliminated on average 78% (p<0.001) or 77% (p=0.006) of the tumor cells, respectively. However, similarly to the findings with untreated control pTc HROC113, the addition of the above-mentioned immune modulators did not result in increased tumor cell killing. Instead, the combination of ICI and PI-9i even seemed to worsen the outcomes with all three T cell populations.

**Figure 5:**
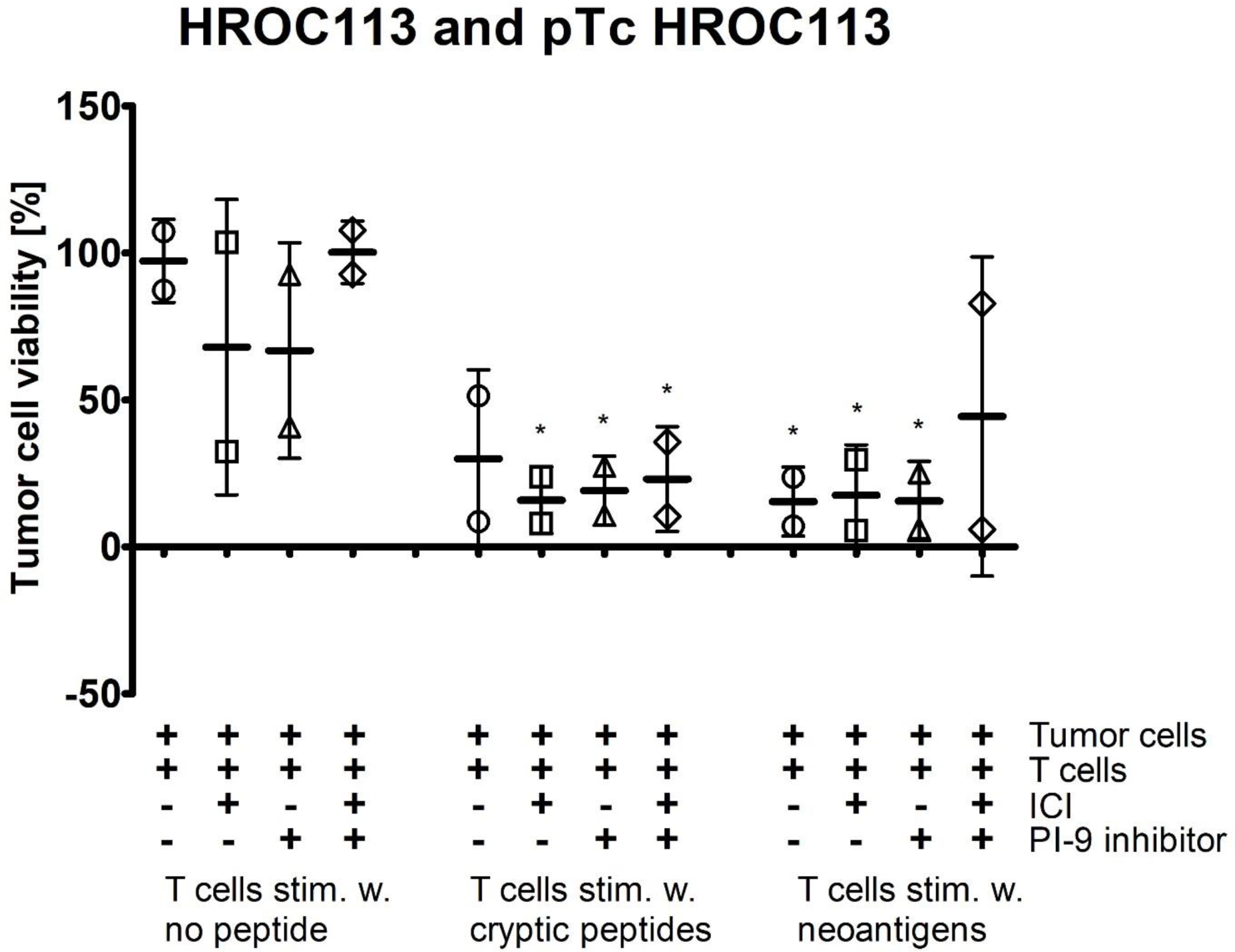
Cytotoxicity of pTc HROC113. Tumor cells were incubated for 48h with pTc HROC113, which were pre-stimulated for 14 days with tumor-specific cryptic or neoantigenic peptides. pTc HROC113 stimulated without peptide served as control. Medium was supplemented with three ICI (20µg/ml pembrolizumab, 10µg/ml ipilimumab and 20µg/ml durvalumab) and/or 400µmol PI-9i. (*) p<0.05 in *t*-test of sample and unstimulated control without immune modulators. Depicted are two biologically independent measurements, mean and standard deviation.

## 3. Discussion

In this study, we investigated the interaction of carefully selected tumor and autologous T cells *in vitro.* In initial co-culture experiments, no anti-tumor responses were observed, which was due to the clinical manifestation of the original tumors rather expected than surprising. Disappointingly, subsequent ICI treatment to re-establish T cell activity and tumor cell recognition did not work out as expected.

Resistance against ICI is either an innate tumor cell property or is observed as an acquired phenomenon after therapy. ICI resistance can be categorized into three groups: resistance mechanisms related to (I) antigen recognition; (II) T cell migration/infiltration; and (III) effector functions of T cells ^21^. When comparing gene expression of proteins involved in antigen processing ^14^ between our two tumor models and normal colon tissue, the majority of genes showed a higher expression in tumor than in normal tissue; hinting at a preservation of functional antigen processing. Even though B2M expression was decreased, a HLA ligandome analysis revealed thousands of HLA-eluted peptides in both cell lines ^20^. Thus, the presentation of tumor-specific peptides with the potential to activate T cells was maintained in these two CRC cases, contrasting the well-established correlation of MSI and diminished HLA expression ^22–24^. However, despite preserved functional tumor antigen presentation, the interaction of the T cell receptor and peptide-presenting tumor cells can be impaired, for example, by an overexpression of cell surface glycosphingolipids sterically impeding the interaction ^25^.

Since antigen presentation was likely not defective and the selected co-culture *in vitro* assay neglected the issue of T cell infiltration, the observed ICI resistance was probably caused by resistance mechanisms related to T cell effector functions. When ICI were used to interfere with PD-1 and PD-L1, which was expressed on the majority of tumor cells, there was no improvement in anti-tumor response observed, even when these ICI where combined with anti-CTLA-4. The detailed characterization of the T cell populations revealed the expression of a further immune checkpoint, namely LAG-3, on the majority of T cells, possibly impairing their activation after binding to tumor cell MHC II. Currently, several anti-LAG-3 antibodies are tested in pre-clinical and clinical trials ^26^. Moreover, as TIM-3 and TIGIT also seem to negatively regulate T cell activity, blocking antibodies are similarly under investigation ^27^ and further immune checkpoints are identified continuously ^28^. By focusing the investigations on clinically approved ICI the results of the study may be limited, but they also prove that further research is urgently needed.

The effector function of T cells is not only suppressed directly by tumor cells, but is mainly influenced by the tumor microenvironment. Here, immunosuppressive cell populations and cytokines are responsible for decreased T cell activity. The determination of regulatory T cell subsets in our selected cancer cases did not indicate strong cell-based immune inhibition as these did not exceed 10%, which equals to regulatory T cell levels in healthy individuals ^29^. Studies claiming increased proportions of regulatory T cells in peripheral blood of tumor patients compared to healthy donors ^30–32^ could thus not be validated in the CRC cases analyzed. Of note, T cells underwent two weeks of unspecific expansion prior to measurements, potentially changing the composition of cell subpopulations. Likely, persisting immune suppression by tumor cells in the tumor microenvironment resulted in lower anti-tumor reactivity of TiTc compared to pTc, characterized by a higher proportion of exhausted T cells or an increased state of T cell exhaustion. Moreover, secretion of known immunosuppressive cytokines like IL-10 could be excluded as driver of T cell inhibition.

Additional examinations revealed granzyme B activity in both tumor cell lines HROC113 and HROC285 T0 M2, accompanied by the expression of its inhibitor PI-9. Granzyme B, which is physiologically expressed in cytotoxic T and natural killer cells ^33^, was already detected previously in breast, lung and urothelial carcinoma cells ^34–36^. Moreover, the expression of PI-9, the protective shield against intracellular granzyme B, was found in prostate, lung and rectal cancer ^37–39^ and we could confirm this finding here for a small series of low-passaged, patient-derived CRC cell lines. Since several studies showed a negative correlation between PI-9 expression and response to ICI ^40,41^, we hypothesized, that the use of PI-9i might restore anti-tumor immune responses.

However, the proof-of-concept cytotoxicity assay of pTc HROC113 and the tumor cell line HROC113 revealed, that neither ICI nor PI-9i treatment resulted in significant changes and even the combination failed to improve tumor cell elimination. It is thus safe to conclude, that tumor cells utilized immunosuppressive mechanisms beyond the investigated cytokine secretion, immune checkpoint expression and adaption to cytotoxic microenvironment. Clinically, failure of ICI would have been interpreted probably as an immunological cold tumor, but T cell stimulations disproved this assumption. When T cells were stimulated with tumor-specific peptides, recognition and elimination of tumor cells increased significantly. Both, cryptic peptides as well as neoantigens, sufficiently induced T cell effector function, clearly exceeding the effects of ICI and/or PI-9i. By comparing stimulated pTc and TiTc we further showed, that T cells from peripheral blood are a valuable source of lymphocytes, which can be turned into tumor-reactive T cells by stimulation with tumor-specific peptides. In connection with studies showing that the quality ^42^ and clonality ^43^ of neoantigens was superior in predicting T cell response in comparison to mere neoantigen number, these results hint at a likely broader applicability of immunotherapy beyond tumors with high TMB.

Nevertheless, the incomplete elimination of tumor cells by peptide-stimulated T cells might again be an indicator of a complex network of immunosuppressive mechanisms in use by the tumor cells, which must be examined further. Our data also imply, that the development of new immune modulators interfering with immunosuppressive molecules (immune checkpoint ligands, enzymes, cytokines, *etc.*) and tumor-individual selection of those will contribute equally to optimally support the effector function of tumor-reactive T cells. By applying *ex vivo* expanded and selected T cells or genetically modified CAR T cells or TCR T cells, the strength of the anti-tumor reactions increases dramatically and can overcome immunosuppressive barriers. Additionally, the combination of strongly primed T cells with further immune modulators could prevent acquired resistance to immunotherapy and tumor relapse as several studies already showed durable responses in combination therapy ^44,45^.

Summarizing all results, our detailed investigation of two distinct sets of patient-derived cell populations demonstrated that ICI-resistant tumor cells can be targeted by T cells primed with tumor-specific antigens. This peptide stimulation induced increased tumor cell recognition not only in TiTc, but also in pTc, which partially even exceeded the former. These findings in pTc could hint towards an equivalent or even superior performance in future immunotherapeutic approaches. In our investigation, extensive T cell priming in a personalized approach was superior to standardized ICI therapy, underlining the importance of precision medicine also in cancer immunotherapy. Additionally, this patient-individual approach could be combined with drugs like ICI and enzyme blockers according to the respective tumor characteristics. Such multimodal approaches could then tackle the complex network of immunosuppressive strategies to successfully overcome immunological barriers erected by any given tumor.

## 4. Material and Methods

### 4.1 Cell Culture

Patient-derived CRC cell lines HROC113 and HROC285 T0 M2 from the HROC collection ^19^ were cultured under standard conditions in DMEM/Ham’s F12 medium supplemented with 10% FCS and 2mM L-glutamine. For several experiments, tumor cells were treated for 48h with 200IU/ml IFNγ (Imukin, Boehringer Ingelheim, Ingelheim am Rhein, Germany). Cell culture reagents were obtained from PAN Biotech, Aidenbach, Germany, unless stated otherwise.

### 4.2 T cell isolation and expansion

Isolation of peripheral blood mononuclear cells and pTc from patientś heparinized blood as well as TiTc from vitally frozen tumor pieces was followed by a rapid Tc expansion protocol (REP) as described recently ^20^.

### 4.3 Co-culture

Co-culture was conducted by adapting a previous protocol ^46^. Briefly, 20,000 IFNγ-treated tumor cells were seeded with 200,000 T cells in a 24-well plate coated with anti-human CD28 (Immunotools, Friesoythe, Germany) in DMEM/Ham’s F12 medium, 10% human AB serum, 2mM L-glutamine, penicillin, streptomycin and amphotericin (T cell medium) supplemented with 300IU/ml IL-2. Every 2-3 days, half of the medium was exchanged. At day seven, cells were harvested and seeded with freshly-prepared IFNγ-treated tumor cells in a ratio of 10:1. The co-culture ended at day 14 and cells were further analyzed. The medium of several experiments was supplemented with ICI (20µg/ml Pembrolizumab (Merck, Darmstadt, Germany); 10µg/ml Ipilimumab (Bristol-Myers Squibb, New York City, NY, USA); or 20µg/ml Durvalumab (AstraZeneca, Cambridge, UK)).

### 4.4 T cell stimulation

Autologous B lymphoid cell lines (B-LCL) were used as antigen presenting cells. For peptide-loading, 3×10^6^ B-LCL were incubated at 37°C with 10µg peptide pools in 1ml serum free medium for 1h. Stimulations were performed in 24-well plates with irradiated (30Gy) peptide-loaded B-LCL added at a ratio of 1:4 to 1×10^6^ expanded Tc in 2ml of T cell medium supplemented with 1xITS solution IV and 300IU/ml IL-2 per well. After seven days of co-culture, Tc were harvested, counted and re-stimulated. T cells stimulated with B-LCL without any peptide served as controls.

### 4.5 Flow cytometry

Tumor and T cells were collected and washed with phosphate-buffered saline (PBS). For extracellular staining, cells were incubated in 100µl staining buffer (PBS, 200mM EDTA, 0.5% bovine serum albumin) with the appropriate volume of antibody for 30min at 4°C protected from light. After centrifugation, the staining solution was discarded and cells were washed again with staining buffer. Stained cells were resuspended in staining buffer and measured promptly. Intracellular staining was performed using the InsideStain Kit (Miltenyi Biotec) according to the manufacturer’s instructions and measured promptly. The following anti-human antibodies were used from Immunotools: CD4-APC, CD8-APC, CD25-FITC, and FoxP3-PE or Biolegend (San Diego, CA, USA): CD107a-FITC, CD152-PE, CD223-APC, CD274-FITC, and CD279-FITC. Samples were measured using a BD FACSCalibur™ and data was analyzed by using FCSalyzer 0.9.21 alpha (Sven Mostböck, Vienna, Austria; https://sourceforge.net/projects/fcsalyzer/).

### 4.6 Degranulation assay

The ability of T cells to recognize tumor cells was tested in the degranulation assay ^46^. A 96-well plate was coated with anti-human CD28 (Immunotools) and 100,000 IFNγ-treated tumor cells were seeded together with 200,000 pTc or TiTc per well in Tc medium containing anti-human CD107a-FITC (Biolegend). After 1h incubation, 1µg/ml Brefeldin A (MedChemExpress, Monmouth Junction, NJ, USA) was added, followed by another 4h incubation. Intracellular flow cytometry staining was conducted as described using CD8-APC (Immunotools) and IFNγ-PE (Biolegend). Samples were measured with gates adjusted to detect CD8^+^/CD107a^+^/IFNγ^+^ cells. To receive values for relative degranulation, percentage of triple-positive Tc was normalized to the triple-positive population in Tc not incubated with tumor cells.

### 4.7 Cytokine detection and FGL1 ELISA

For cytokine detection, supernatants were collected from Tc, tumor cells and co-cultures, all cultured for seven days without medium exchange. The concentrations of granulocyte-macrophage colony-stimulating factor (GM-CSF), IFNα, IFNγ, lL-2, IL-4, IL-5, IL-6, IL-9, IL-10, IL-12p70, IL-17A, and tumor necrosis factor (TNF) were determined by MACSPlex Cytokine 12 Kit (Miltenyi Biotec). Supernatants of tumor cell lines were also used for the measurement of FGL-1 with the Human FGL1/Fibrinogen Like Protein 1 ELISA Kit (Assay Genie, Dublin, Ireland).

### 4.8 GranToxiLux

The GranToxiLux assay was performed according to manufacturer’s instructions (Oncoimmunin Inc, Gaithersburg, MD, USA).

### 4.9 Cytotoxicity assay

Tumor cells were seeded into a 48 well plate and stimulated with 200IU/ml IFNγ for 48h. Simultaneously, respective wells were treated with 400µM PI-9i (1,3-Benzoxazole-6-carboxylic acid, Advanced ChemBlocks Inc, Hayward, CA, USA). Following pre-treatment, 5×10^5^ peptide-stimulated or control Tc were added to the approximately 1×10^5^ tumor cells in each well. The number of tumor cells was estimated using previously determined doubling times. The co-culture medium was supplemented with ICI (20µg/ml Pembrolizumab; 10µg/ml Ipilimumab; 20µg/ml Durvalumab) and/or PI-9i (400µM). After 48h of co-culture, medium was discarded, wells were washed once with PBS before 200µl of 0.2% crystal violet solution was added. After 20min incubation at room temperature, staining solution was discarded and wells were washed three times with PBS. When plates were dry, 500µl of sodium dodecyl sulfate solution was added and optical density at 570nm was measured at the microplate reader Infinite 200 (Tecan, Männedorf, Switzerland).

### 4.10 RNA expression analysis

RNA sequencing of the patient-derived cell lines was performed by the Center for Quantitative Biology (QBiC, University of Tübingen, Germany) and resulting data was provided. Moreover, publicly available data was used for comparison. First, R package (R (4.2.1) version) was applied to The Cancer Genome Atlas (TCGA, https://portal.gdc.cancer.gov/ (accessed on 20.06.2022)) and the Genotype-Tissue Expression (GTEx, gtexportal.org/home/ (accessed on 20.06.2022)) databases to analyze the difference of the mRNA levels of 18 genes of interest including ERAP1 [ENSG00000164307.12], ERAP2 [ENSG00000164308.16], PDIA3 [ENSG00000167004.12], PSMB5 [ENSG00000100804.18], PSMB6 [ENSG0000142507.9], PSMB7 [ENSG00000136930.12], PSMB8 [ENSG00000204264.8], PSMB9 [ENSSG00000240065.7], PSMB10 [ENSG00000205220.11], B2M [ENSG00000166710.17], CALR [ENSG00000179218.13], CANX [ENSG00000127022.14], TAP1 [ENSG00000168394.10], TAP2 [ENSG00000204267.13], TAPBP [ENSSG00000231925.11], HLA-A [ENSG00000206503.11], HLA-B [ENSG00000234745.9] and HLA-C [ENSG00000204525.14]. Gene expression was compared between 484 CRC and 359 normal colorectal mucosal tissue samples. Moreover, TMB was used to group the cancer samples: Samples were sorted by TMB and top 20% were identified as TMB high. Remaining samples were considered TMB low.

### 4.11 Statistical analyses

Statistical significance was determined by an unpaired, two-sided *t*-test using GraphPad Prism 5 (Boston, MA, USA). Results were stated significant if the *t*-test resulted in p value <0.05. Details on compared groups are given in the figure legends. If not stated otherwise, graphs depicting the results show mean values and the respective standard deviation.

## Ethics approval and consent to participate

The study was conducted according to the guidelines of the Declaration of Helsinki. All patients gave informed written consent to participate in the study and all procedures were approved by the Ethics Committee of the University of Rostock University Medical Center (Reference numbers: A 2018-0054 and A 2019-0187) in accordance with generally accepted guidelines for the use of human material.

## Acknowledgements

The authors express their gratitude to the patients for participating in the HROC collection. This research was in part funded by grant number TBI-V-1-241-VBW-084 and TBI-V-240-VBW-084 from the state Mecklenburg-Vorpommern.

## Author Contributions

Design and conceptualization of the study was directed by ML. Experimental design, data interpretation and formal analysis was performed by SS. MK established the patient-derived cell lines and SZ analyzed expression data. Identification and validation of cryptic peptides and neoantigens was performed by MWL and AS. The initial draft and figures were prepared by SS and ML; SZ, MK, MWL and AS reviewed and edited the manuscript. All authors have read and agreed to the final version of the manuscript.

## Data Availability

The datasets analyzed during the current study available from the corresponding author on reasonable request.

