## Supplementary figures for "Peptide-stimulation of autologous T cells reverts immune checkpoint inhibitor resistance of hypermutated colorectal cancer cells"

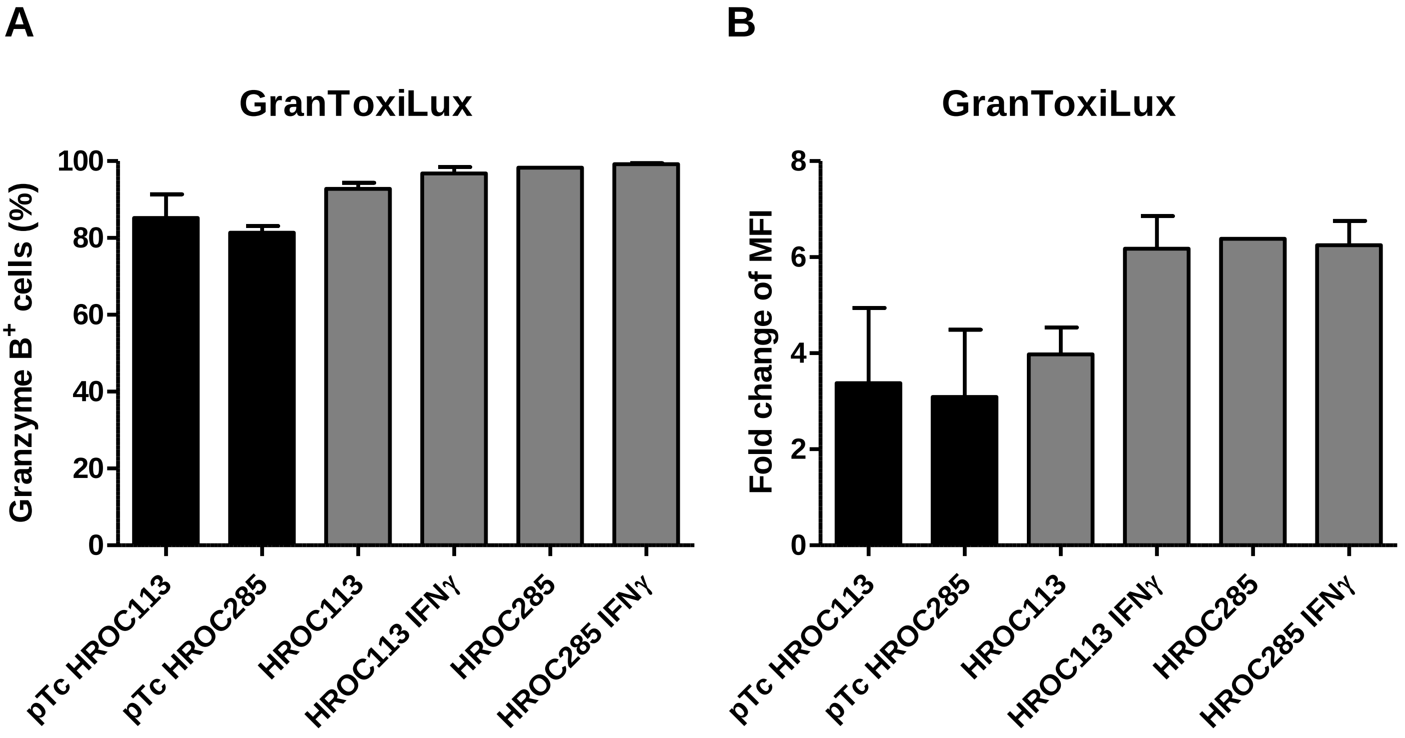


**Figure S1: Granzyme B activity in CRC cell lines.** The GranToxiLux kit (Oncoimmunin Inc, Gaithersburg, Maryland, USA) was used to measure the granzyme B activity in tumor cells and pTc, which were used as positive controls. The amount of cells with granzyme B cleaving capacity was determined (A) as well as the MFI of granzyme B compared to unstained controls (B). Depicted are means of 2-3 biological replicates and the respective standard deviation.

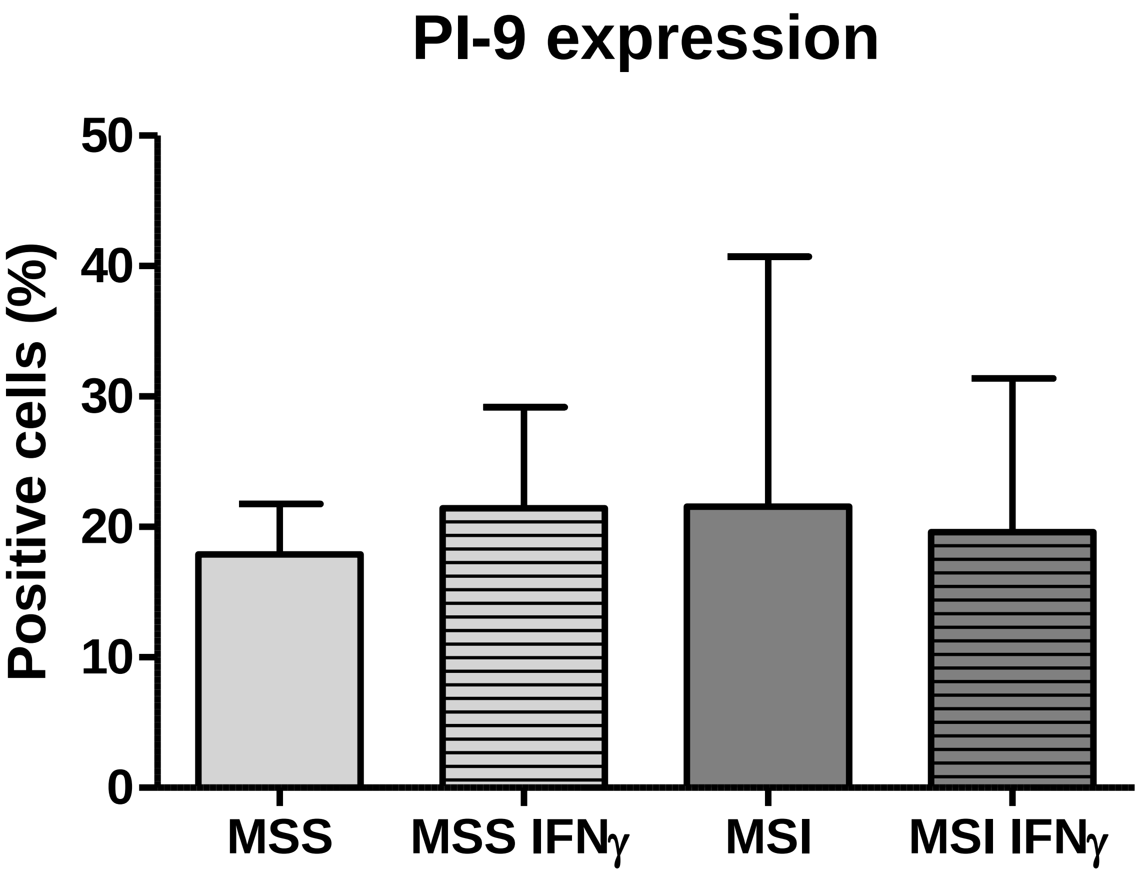


**Figure S2: PI-9 expression in microsatellite stable and unstable CRC cell lines.** Comparison of percent PI-9$^+$ cells between microsatellite stable (MSS; n=4) and MSI (n=5) cell lines determined by flow cytometry. Depicted are means of the biological replicates and the respective standard deviation.

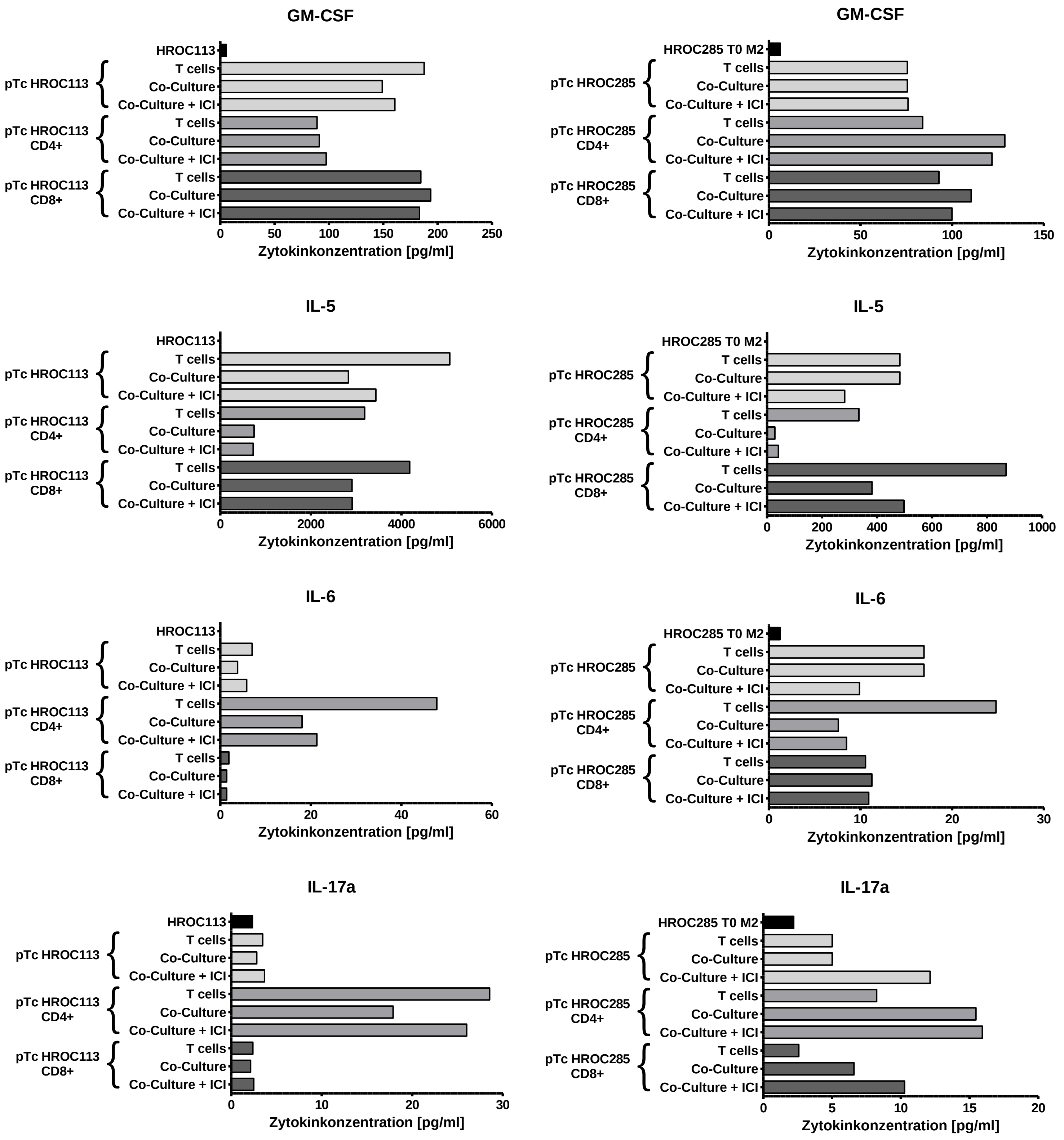


**Figure S3: Cytokine detection in cell culture supernatants of tumor cell lines.** Cell culture supernatants were collected after seven days of culture or co-culture of pTc and their respective tumor cells. Cytokine concentration was determined by using the MACSPlex Cytokine 12 Kit from Miltenyi Biotec. For IL-10, IL-4, -9, -10, -12p70, IFNα and TNF, samples did not reach the limit of detection. Depicted are mean values of two technical replicates.

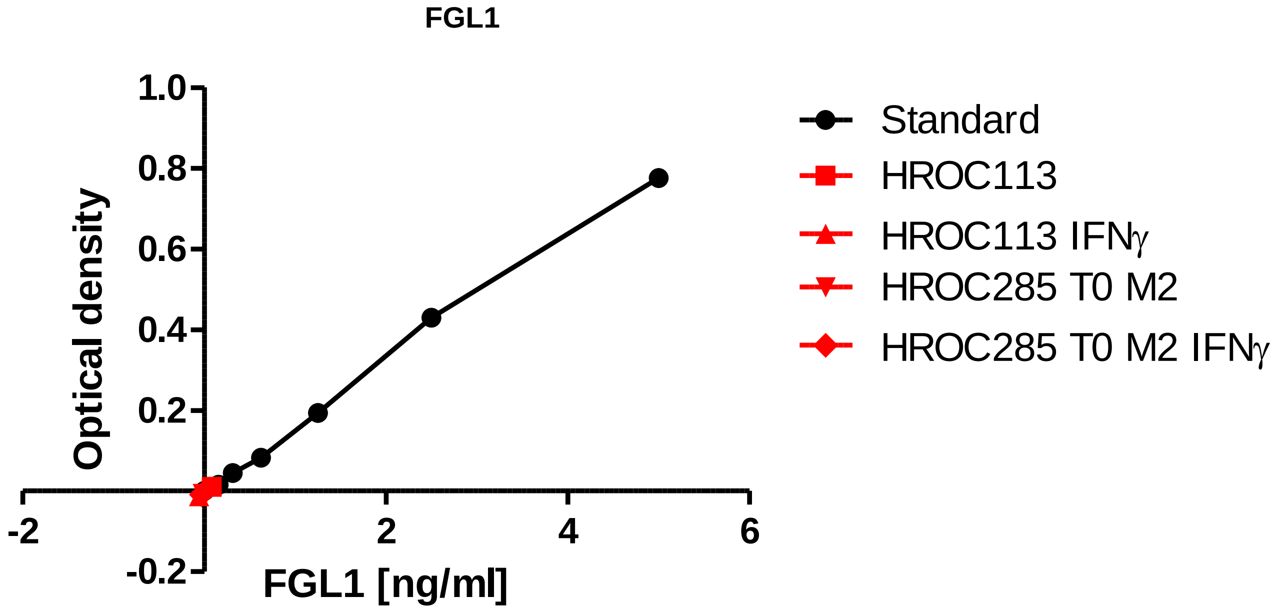


**Figure S4: FGL1 secretion by tumor cells.** Tumor cells were cultivated with or without 200 IU/ml IFNγ and supernatant was collected at day four when cell growth reached confluency.

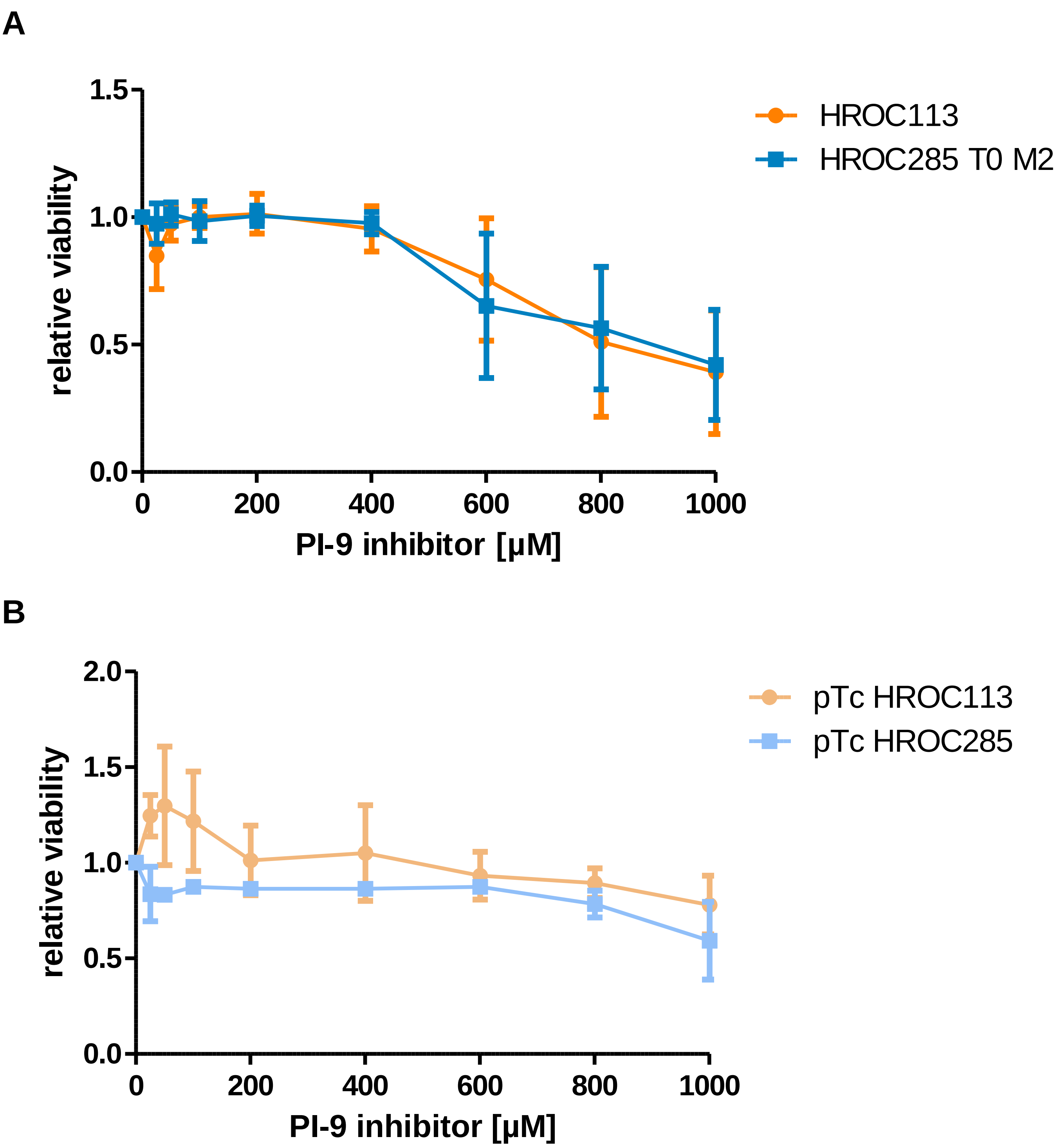


**Figure S5: Effect of PI-9 inhibitor on cells.** Tumor cells (A) and pTc (B) were incubated with increasing concentrations of the PI-9 inhibitor 1,3-Benzoxazole-6-carboxylic acid for seven days. Subsequently, viability of tumor cells and T cells was determined by crystal violet staining and calcein AM assay, respectively.
